# Exponentiating pixel values for data augmentation to improve deep learning image classification in chest X-rays

**DOI:** 10.1101/2021.03.11.434925

**Authors:** Takuma Usuzaki, Kengo Takahashi, Daiki Shimokawa, Kiichi Shibuya

## Introduction

Data augmentation (DA) is a technique of artificial intelligence (AI) and mainly used with convolutional neural network (CNN) to enhance the size and quality of datasets.^1^ CNN with DA can improve prediction accuracy of Chest X-ray (CXR) screening and lead to reducing burden on radiologists.^2,3^ A previous study reported that exponentiating pixel values of a medical can be used for DA through enhancing contrast of the image.^4^ However, it is still unclear whether exponentiating pixel values of CXR images can be used for DA with CNN. The aim of this study is to evaluate the effectiveness of exponentiating pixel values for DA using CNN in the task of classifying normal and abnormal CXR images.

## Method

“Japanese Society of Radiological Technology, chest X-ray image dataset” that includes 93 normal images and 154 abnormal images with a lung nodule (2048 × 2048 matrix, 0.175-mm pixel size), was used^5^. We randomly divided 154 normal images and 93 abnormal images into training (92 normal, 55 abnormal) and test (62 normal, 38 abnormal) dataset. In constructing an image for DA, each pixel value of the original images was exponentiated by the exponent ranging from 1 to 10, incrementing by 0.5. We call this image as exponentiated image (EI). For each exponent (1.0, 1.5, 1.5, 2.0, …, 10.0), the CNN model was trained using the original training dataset and EI for 40 epochs. Test accuracy was calculated at the end of each epoch using the test dataset. The maximum test accuracy (MTA) among the 40 test accuracies was saved for statistical analysis. This process was repeated 50 times for each exponent (1.0, 1.5, 1.5, 2.0, …, 10.0). The mean MTA for each exponent (1.5, 1.5, 2.0, …, 10.0) was compared using Student’s t-test, to that of 1.0. A Visual Geometry Group 16 (VGG16) model trained by ImageNet was used by resizing all images to 224 × 224 and normalizing.^6^ During the training process, binary cross entropy was optimized with Adam optimizer (learning rate=0.0001, β1=0.9, β2=0.999, ε=1.0×10^−8^, weight decay=0, AMSGrad=False). All analyses were performed using Python Language, version 3.8.2 (Python Software Foundation at http://www.python.org). Statistical significance was defined as p-value<0.05.

## Result

The mean MTA when the exponent was 1.0 was 0.749 (reference). The mean MTA was higher than the reference at exponent values 4.5(MTA=0.762, p-value=0.014), 5.0(0.762, 0.019), 5.5(0.772, 2.9×10^−5^), and 6.5(0.763, 0.010). Figure shows the means±standard deviations (SD) of MTA for each exponent (1.0, 1.5, 2.0, …, 10.0). Table shows the mean MTA, SD of MTA, and the p-value of Student’s t-test for each exponent (1.0, 1.5, 2.0, …, 10.0).

**Figure 1.**
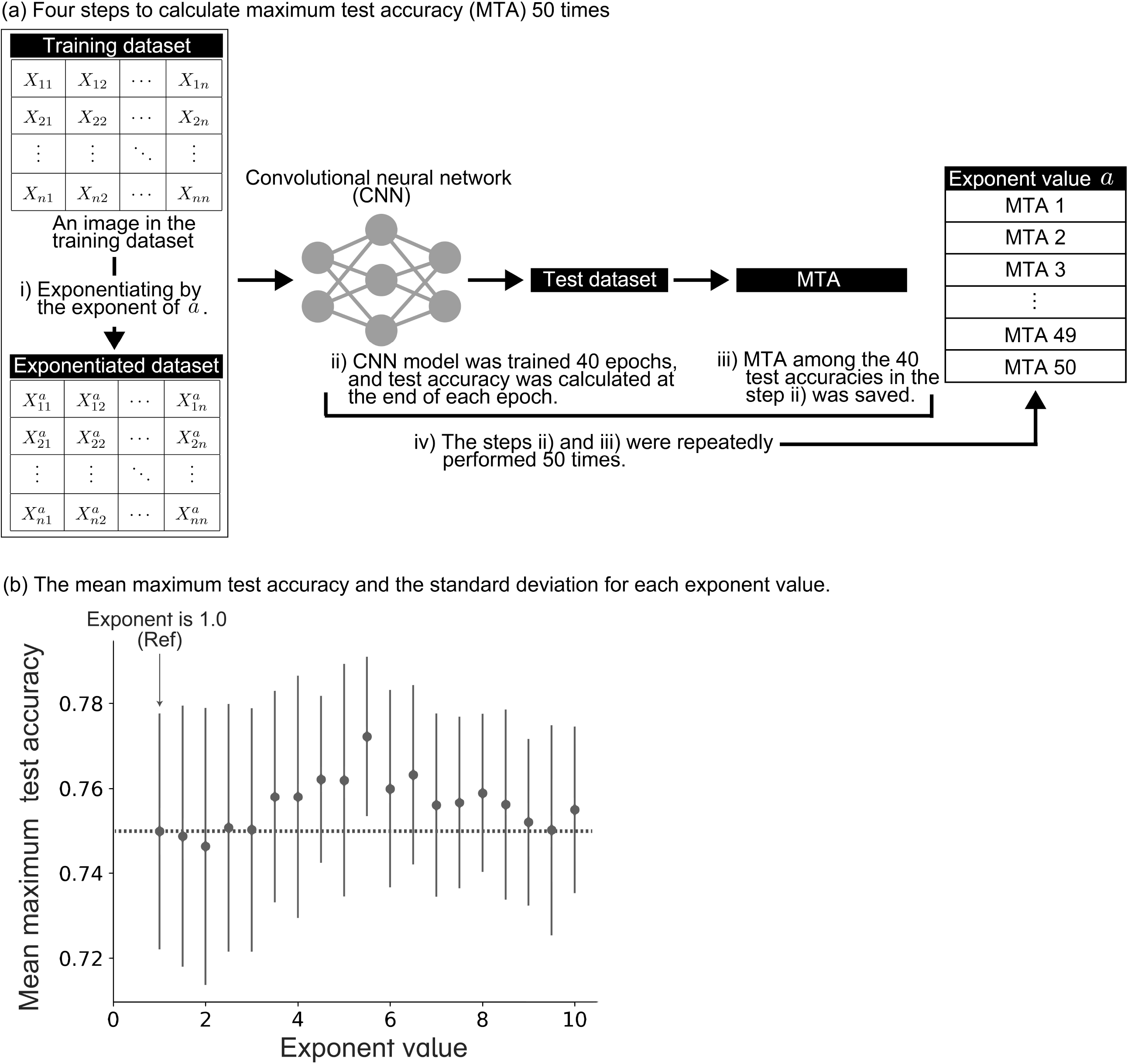
Maximum test accuracy for each exponent. Means±standard deviations of maximum test accuracy (MTA) for each exponent (1.0, 1.5, 2.0, …, 10.0). The horizontal and vertical axes represent the exponent value and MTA, respectively. The arrow points to the case the exponent value is 1.0, which is used as the reference value. The broken line represents the reference value of the MTA.

## Discussion

In our analysis, accuracy of CNN trained by EI was improved when classifying normal and abnormal CXR images. This result indicates exponentiating pixel values can be used for DA. An EI has extended pixel values which are partially outside of the gray scale range. This extension of pixel values may improve the CNN performance through contrast enhancement.^4^

Our study has two limitations. First, the number of images we examined were limited. The effectiveness of exponentiating pixel values for DA should be evaluated on a larger scale. Second, abnormal images were not classified by diseases. An accuracy change depending on exponent should be examined to each disease because the dependence may differ in each disease. The accuracy change depending on exponent may be used to classify diseases in clinical practice. However, our results indicate that CNN trained by EI improve accuracy in the CXR screening process. Further studies are needed to clarify the properties of exponentiating for DA.

## Conclusion

Exponentiating pixel values can be used for DA with CNN to classify normal and abnormal CXR images.

## Notes

### Competing Interest Statement

The authors have declared no competing interest.

