## Supplementary material for "Exponentiating pixel values for data augmentation to improve deep learning image classification in chest X-rays": Table

Table 1

The mean maximum test accuracy (MTA), standard deviation (SD) of MTA and the p-value of Student's t-test for each exponent.

| Exponent | 1.0 (Ref) | 1.5 | 2.0 | 2.5 | 3.0 | 3.5 | 4.0 | 4.5 | 5.0 |
|---|---|---|---|---|---|---|---|---|---|
| Mean MTA | 0.750 | 0.749 | 0.746 | 0.751 | 0.750 | 0.758 | 0.758 | 0.762 | 0.762 |
| SD | 0.0278 | 0.0308 | 0.0327 | 0.0292 | 0.0287 | 0.0249 | 0.0286 | 0.0197 | 0.0274 |
| p-value | | 0.84 | 0.55 | 0.88 | 0.94 | 0.13 | 0.18 | 0.014 | 0.019 |

| Exponent | 5.5 | 6.0 | 6.5 | 7.0 | 7.5 | 8.0 | 8.5 | 9.0 | 9.5 | 10.0 |
|---|---|---|---|---|---|---|---|---|---|---|
| Mean MTA | 0.772 | 0.760 | 0.763 | 0.756 | 0.757 | 0.759 | 0.756 | 0.752 | 0.750 | 0.755 |
| SD | 0.0188 | 0.0233 | 0.0212 | 0.0216 | 0.0202 | 0.0186 | 0.0224 | 0.0196 | 0.0248 | 0.0197 |
| p-value | $2.9 \times 10^{-5}$ | 0.095 | 0.010 | 0.21 | 0.077 | 0.081 | 0.24 | 0.65 | 0.95 | 0.31 |

Abbreviations: Ref, reference; MTA, maximum test accuracy; SD, standard deviation.

p-value < 0.05 columns are gray colored.
